# Anchor negatively regulates BMP signaling to control Drosophila wing development

**DOI:** 10.1101/052217

**Authors:** Xiaochun Wang, Ziguang Liu, Li hua Jin

## Abstract

**Summary statement:** The novel gene *anchor* is the ortholog of vertebrate GPR155, which contributes to preventing wing disc tissue overgrowth and limiting the phosphorylation of Mad in presumptive veins during the pupal stage.

G protein-coupled receptors play a particularly important function in many organisms. The novel *Drosophila* gene *anchor* is the ortholog of vertebrate GPR155, and its molecular function and biological process are not yet known, especially in wing development. Knocking down *anchor* resulted in increased wing size and extra and thickened veins. These abnormal wing phenotypes are similar to those observed in gain-of-function of BMP signaling experiments. We observed that the BMP signaling indicator p-Mad was significantly increased in *anchor* RNAi-induced wing discs in larvae and that it also abnormally accumulated in intervein regions in pupae. Furthermore, the expression of BMP signaling pathway target genes were examined using a *lacZ* reporter, and the results indicated that *omb* and *sal* were substantially increased in *anchor* knockdown wing discs. In a study of genetic interactions between Anchor and BMP signaling pathway, the broadened and ectopic vein tissues were rescued by knocking down BMP levels. The results suggested that the function of Anchor is to negatively regulate BMP signaling during wing development and vein formation, and that Anchor targets or works upstream of Dpp.

## INTRODUCTION

*Drosophila*, as a practical model, is used to study the wing pattern formation. Cells within wing discs must receive many different signals to form an appropriate pattern. This variety of signals controls an enormous array of cellular processes, such as proliferation, differentiation, apoptosis, and cell migration (Neufeld, et al., 1998; Egoz-Matia, et al., 2011; Shbailat and Abouheif, 2013). The processes that control vein formation during wing development are, in particular, mediated by many different signaling pathways, including the Bone Morphogenetic Protein (BMP), Epidermal Growth Factor (EGF), Hedgehog, Notch and Wnt pathways (Blair, 2007).

A BMP gradient is an important signaling mechanisms that is required for wing patterning in *Drosophila*. BMPs are members of the transforming growth factor-β (TGF-β) subfamily, the involvement of which is conserved in many patterning processes across multicellular organisms (Clarke and Liu, 2008). In *Drosophila*, the BMP family includes two classical ligands and three receptors. The ligands are the BMP2/4 homolog Decapentaplegic (Dpp) and the BMP5/6/7/8 homolog Glass bottom boat (Gbb) (Garcia-Bellido and Merriam, 1971; Wharton et al., 1991). The receptors include two type I receptors, Thick-veins (Tkv) and Sax (Saxophone), and a type II receptor, Punt (Put) (Affolter et al., 1994; Xie et al., 1994; Letsou et al., 1995). Dpp provides a long range signal along the anterior-posterior (AP) boundary of the wing disc, while Gbb provides a longer range signal than Dpp in the wing disc (Ray and Wharton, 2001). Dpp and Gbb form homo-or heterogeneous dimers that elicit effects during wing development (Israel et al., 1996). When a ligand binds to a receptor on the surface of cell, the BMP family R-Smad member Smad 1/5/8 and the intracellular Smads (R-Smads) are phosphorylated, allowing them to recruit Co-Smad (Smad 4, Medea) and then translocate into the nucleus, where they regulate the transcription of their target genes (Chacko et al., 2001; Shi and Massagué, 2003). BMP’s target genes include *optomotor blind* (*omb*) (Sivasankaran et al., 2000), *spalt* (*sal*) (Barrio and de Celis, 2004), *daughter against dpp* (*dad*) (Tsuneizumi et al.1997), and *brinker* (*brk*). In particularly, *brk* exprssion is repressed by Dpp (Campbell and Tomlinson, 1999). These genes can affect cells fate decisions and tissue patterns to varying degrees. Dpp is a typical factor that is involved in all currently characterized BMP signaling events in *Drosophila*, and it performs developmental and genetic functions in the wing disc. During larval development, Dpp is expressed in a strip along the AP boundary to promote wing disc proliferation and specify presumptive vein formation (Capdevila and Guerrero, 1994; Biehs et al., 1998). Knocking down Dpp signaling lead to the development of small discs, reduced adult wing size and resulted in the loss of veins (Spencer et al., 1982; Künnapuu et al., 2009). During the pupal stage, Dpp is localized at the AP boundary and along presumptive veins. *dpp* mutants have the shortveins phenotype, indicating that Dpp promotes the formation of distinct intervein patterns (Segal and Gelbart, 1985). In addition, the ectopic activation of Dpp signaling induces the overgrowth of wing tissues (Martín-Castellanos and Edgar, 2002; O’Keefe et al., 2014). *gbb* is broadly expressed in the wing disc except at the AP boundary strip, and *gbb* mutants exhibit phenotypes similar to those of *dpp* mutants, have small wing disc and are missing veins (Khalsa et al., 1998; Ray and Wharton, 2001).

Unlike typical GPCRs, the mouse GPR155 sequence predicts that it has 17 transmembrane regions. Five variants of GPR155 mRNA have been identified, including the Variant 1 and Variant 5 proteins, which have an intracellular DEP domain near the C-terminal (Trifonov et al, 2010). GPR155 has only been reported in mouse, and evidence suggests that GPR155 plays specific roles in Huntington disease and autism spectrum disorders (Brochier et al., 2008). However, the molecular function and biological processes that are affected by *Drosophila* GPR155 have not been reported, including in wing development. *Drosophila* CG7510, which we have called *anchor*, is an integral membrane protein that is the ortholog if vertebrate GPR155, and it shares more than 37% homology with the mammalian GPR155. In this study, we knocked down *anchor* using wing-specific Gal4 lines, and we observed enlarged wing size, and a thickened and ectopic veins phenotype. During larval and pupal stages, we detected ectopic levels of p-Mad, which is downstream of BMP signaling, when *anchor* RNAi was applied. This also increased the levels of target genes such as *sal* and *omb*. Taken together, those results suggest that the novel functions of Anchor are mediated through the negative regulation of BMP signaling to control wing development in *Drosophila*.

## RESULTS

### Locus of the *anchor* gene and homolog analysis

The transcription site of *anchor* is located at 74E2-74E3 on chromosome 3 in *Drosophila*, and its open reading frame (ORF) encodes a predicted protein that is 949 amino acid residues long (Fig. 1A). A membrane protein called Anchor was identified in a systematic search for genes that are homologous to known G-protein coupled receptors (GPCRs). The amino acid sequence of the Anchor gene contains 17 transmembrane domains and a C-terminal DEP domain (Fig. 1B). *anchor* encodes a protein with more than 37% homology to GPR155 in humans, chimpanzees, mice and rats (Fig. 1C). The conserved sequence in the DEP domain was first observed in three proteins: Dishevelled (*D. melanogaster*), which is an adaptor of the Wingless (Wnt) signaling pathway (Klingensmith et al., 1994); EGL-10 (*C. elegans*), which is a negative regulator of GPCR signaling (Koelle and Horvitz, 1996); and mammalian Pleckstrin, which mediates signaling in platelets and neutrophils (Kharrat et al., 1998).

**Fig. 1.**
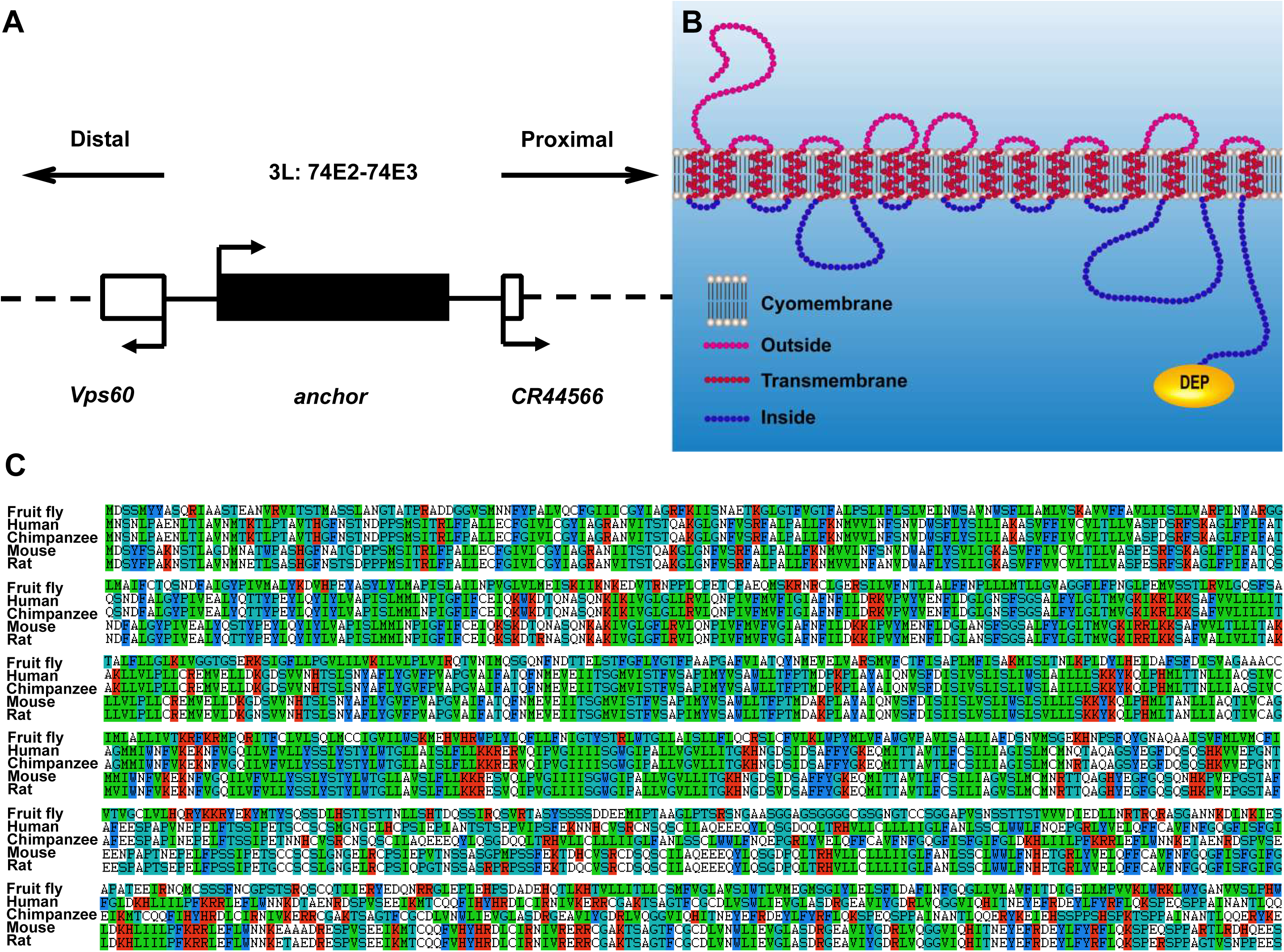
Molecular characteristics of *Drosophila* Anchor. (A) The transcriptional unit containing *anchor* is located at 74E2-74E3 on chromosome 3. (B) Diagram of the predicted secondary structure of Anchor. Potential 17-transmembrane domains and a DEP domain were identified using TMHMM software (http://www.cbs.dtu.dk/services/TMHMM/). The transmembrane domain amino acids are indicated by red lines. (C) Protein alignment was performed using Clustal X with the following sequences: fruit fly (*Drosophila melanogaster*, NP_648998.2), human (*Homo sapiens*, NP_001253979.1), chimpanzee (*Pan troglodytes*, XP_003309468.1), mouse (*Mus musculus*, NP_001177226.1), and rat (*Rattus norvegicus*, NP_001101281.1). The fruit fly *anchor* shared 37% homology with the human sequence, 37% with the chimpanzee sequence, 38% with the mouse sequence, and 38% with the rat sequence.

### Knocking down *anchor* induced extra vein formation in adult flies

To study the *in vivo* role of Anchor in *Drosophila* development, we used an *anchor* RNAi attached to a ubiquitously expressed promoter (e.g., *Actin-Gal4* and *Tubulin-Gal4*). Unfortunately, the RNAi induced a severe embryonic lethal phenotype. Next, we combined the *anchor* RNAi with *Hs-Gal4* at 25°C and then shifted the temperature to 37°C for 90 min each day beginning in the third instar larval stage and continuing until the flies were adults. Surprisingly, we found that the adult wings of these flies exhibited abnormal wing veins (data not shown). These results led us to speculate that the novel gene *anchor* plays an important role in *Drosophila* wing development. To determine whether the abnormal wing phenotypes were caused by the autonomous loss of the Anchor protein, we used *A9-Gal4* and *MS1096-Gal4* to ubiquitously knockdown *anchor* in the wing blade. Unexpectedly, these flies displayed thicker and more veins than the wings in the control flies (Fig. 2A-B, H-I); When we used the *dpp-Gal4*, *ptc-Gal4* and *en-Gal4* drivers, we also observed ectopic vein tissues in the *anchor* RNAi-induced adult wings (Fig. 2C-E, J-L), and we observed that the extra veins in these flies were restricted to the AP boundary and the anterior or posterior compartment of the wing. *ap-Gal4* is specifically expressed in the dorsal cells of the L3 wing disc, while *brk-Gal4* is expressed in the tissues surrounding the wing pouch cells in the L3 wing disc and the intervein cells in pupal wings (Campbell and Tomlinson, 1999; Sotillos and De Celis, 2005). As expected, we observed wing expansion defects when we used the *ap-Gal4* and *brk-Gal4* drivers. In addition, a blistering phenotype was observed in *ap>anchor* RNAi wings (Fig. 2 F-G, M-N).

**Fig. 2.**
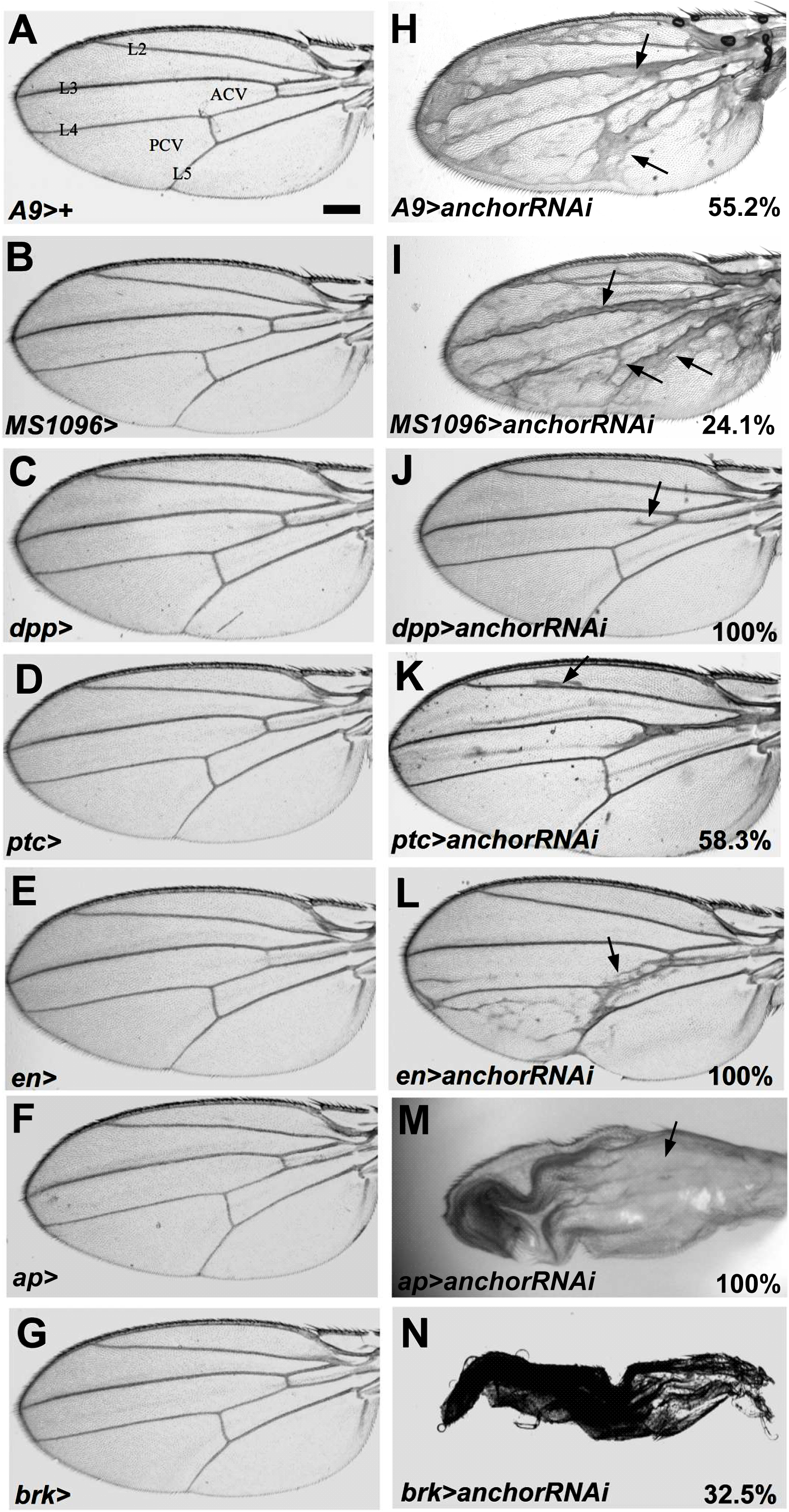
Wing-specific knockdown of *anchor* resulted in thickened and extra veins in adult flies. (A-G) Control wing of a female *Gal4*/*w*^*1118*^ (*Gal4>+*) fly. The wing-specific *Gal4* driver included *A9-Gal4*, *MS1096-Gal4*, *dpp-Gal4*, *ap-Gal4*, *ptc-Gal4*, *brinker-Gal4*, and *en-Gal4*. Longitudinal veins (L2-L5) and crossing veins (ACV, PCV) are indicated in A. (H-N) The *anchor* RNAi flies were crossed with flies carrying a wing-specific *Gal4* driver. These flies displayed wing blades with reduced sizes in differing compartments. (H) The *A9>anchor* RNAi, (I) *MS1096>anchor* RNAi, (J) *dpp>anchor* RNAi, (K) *ptc>anchor* RNAi, and (L) *en>anchor* RNAi resulted in the formation of ectopic and thickened veins. (M) The *ap>anchor* RNAi resulted in wing blistering (arrow). (N) The *brk>anchor* RNAi resulted in reduced wing size and patterning defects. All images were acquired at the same magnification, and all crosses were performed at 29°C (A-C, H-J) or 25°C (D-G, K-N). Scale bar: 200 μm.

We also observed a moderate but consistent increase in wing size in the *A9>anchor* RNAi flies, in which the wings were more than 17% larger than those in the controls (Fig. S1A, C). The distance between L3 and L4 was increased by approximately 29% in the *ptc>anchor* RNAi wings (Fig. S1B, D). In addition, we used another *anchor* RNAi construct (v105969) in combination with the *MS10986-Gal4* and *A9-Gal4* drivers, and we observed that extra veins developed and that wing size was increased (Fig. S2). To provide further confirmation that Anchor is involved in wing development, we constructed transgenic flies in which the entire *anchor* coding region was ectopically expressed. We then combined these flies with *A9>anchor* RNAi flies. As expected, the extra vein phenotype was clearly rescued in flies carrying a functional full-length *anchor* (Fig. S3A, B). *anchor*^*P*^ is a *P*-element insertion in the 5’ UTR region of *anchor* that encodes a Gal4-responsive enhancer. Similar to previous results, we observed that overexpressing *anchor*^*P*^ using the *A9-Gal4* driver as the background for *anchor* RNAi resulted in the rescue of the thickened wing vein phenotype (Fig. S3C, D). These observations suggest that Anchor plays an important role in wing development and especially in wing vein formation. Furthermore, the phenotype of increased vein tissues in the *anchor* RNAi wings is similar to that observed in gain-of-function BMP signaling experiments (Martín-Castellanos and Edgar, 2002; O’Keefe et al., 2014). We therefore proposed that Anchor may be involved in BMP signaling during *Drosophila* wing development.

### The control of Anchor wing size was mediated by an increase in cell proliferation and size in imaginal discs

To determine the biological function of Anchor during wing development, we first analyzed the expression pattern of *anchor* using *in situ* hybridization. We observed that *anchor* is expressed in a generalized manner in imaginal discs (Fig. S4A, C). However, we observed significantly reduced *anchor* levels in RNAi flies that also expressed *MS1096-Gal4* and *A9-Gal4* (Fig. S4B, D). This result further confirmed that the increased wing vein phenotype resulted from low expression levels of the *anchor* gene. We also observed that imaginal discs were enlarged in *anchor* RNAi flies, which led us to speculate that Anchor is required to inhibit excessive cell proliferation in imaginal discs. Wings cell proliferation begins during the larvae stage, and the size of the adult wing is predetermined by the final size of the wing imaginal disc (Day and Lawrence, 2000). The enlarged wing size observed in the *anchor* RNAi flies may have resulted from an increase in cell number or cell size. To explore this, we first analyzed cell proliferation using phospho histone H3 antibodies (PH3), which stain dividing cells that are in M phase. The numbers of PH3^+^ cells were significant higher, by 42%, in the *A9>anchor* RNAi wing discs than in the controls (Fig. 3A, B, G). We further identified the L3 wing blade using Wg antibodies and found that the size of the wing blade in *A9>anchor* RNAi flies was significantly increased by 40% (Fig. 3A’, B’, H). Unexpectedly, we also found that Wg expression levels were significantly increased (Fig. 3A’, B’, I). Next, we compared cell size using phalloidin staining and found that the average size of cells in the *A9>anchor* RNAi flies was higher, by 32%, than the cell size in the controls (Fig. 3D, E, J). To validate our proposal that Anchor is involved in controlling wing and cell size, we overexpressed the full-length *anchor* gene in combination with *A9>anchor* RNAi in flies. As expected, this combination completely rescued the phenotype, including the number of PH^+^ cells, cell size, wing blade and Wg expression levels. Wing size was, in particular, completely rescued (Fig. 3C, C’, F and G-J).

**Fig. 3.**
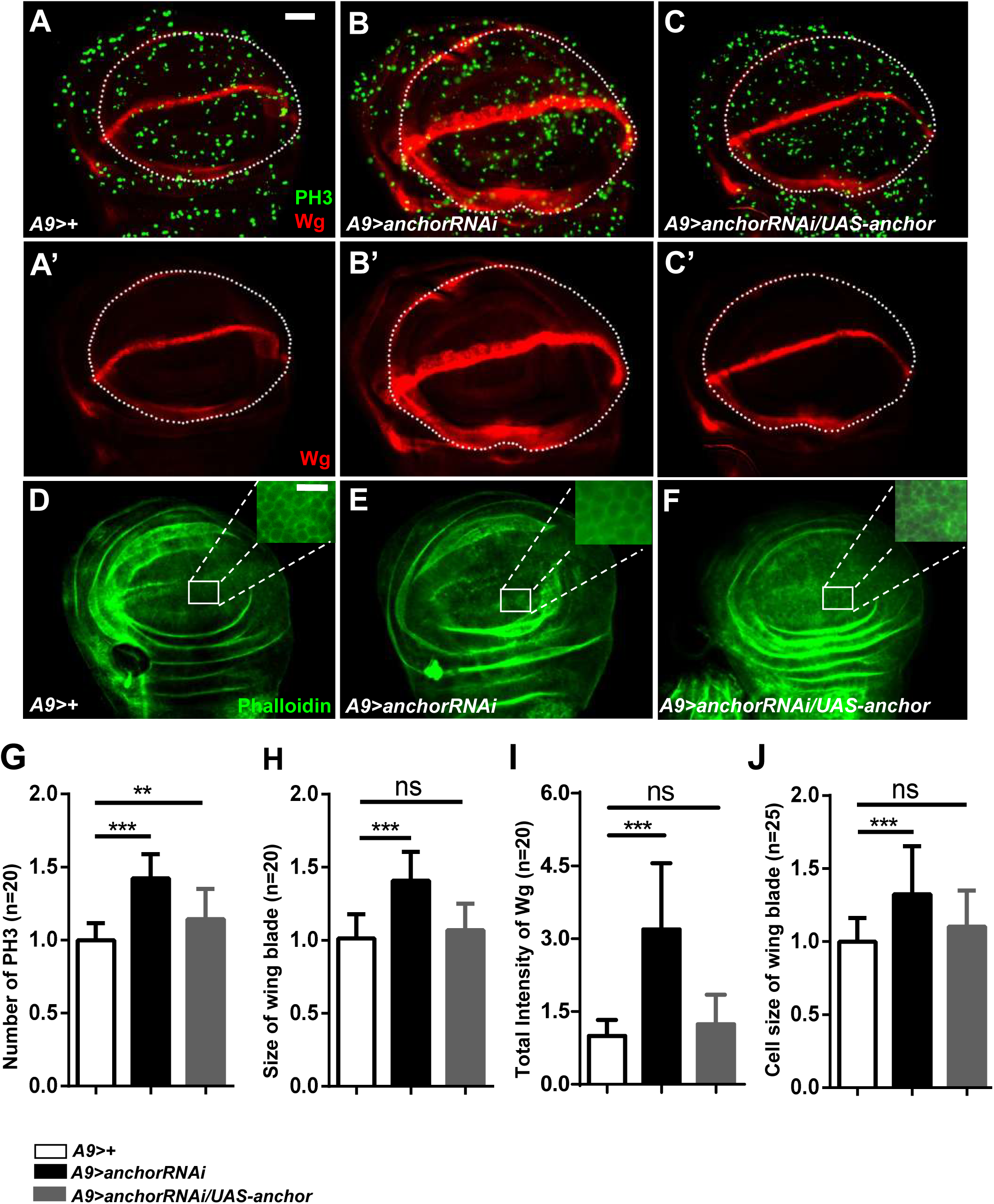
Analysis of cell proliferation and cell size in *anchor* RNAi imaginal discs. (A-C’) Analysis of mitosis and wing blade development in *A9>+*, *A9>anchor* RNAi and *A9>anchor RNAi/UAS-anchor*-induced flies. Cell proliferation was monitored using PH3 (green), wing blade morphology was detected using Wingless (Wg, red), and the wing blade is outlined by a white dotted line. The PH3^+^ cells in the area containing the wing blade and wing blade sizes were significantly increased in *A9>anchor* RNAi-induced flies (B-B’). This phenotype was completely rescued by *UAS-anchor* (C-C’). (D-F) Cell outlines were visualized using phalloidin (green), the white squares in the discs are shown at a higher magnification (Scale bar =10μm), and cell size was increased in the *anchor* RNAi flies. (G-J) Quantification of PH3^+^ cell numbers, wing blade size, Wg labeling intensity and the average size of a single cell in the dorsal area of the wing pouches shown in A-F. All combinations were grown at 29°C and a select group of female larvae were dissected. Scale bar: 50μm

Next, we analyzed apoptopic cell death using a direct TUNEL labeling essay. However, we did not observe any difference between the *anchor* knockdown and control wing discs (data not shown), indicating that the hyperproliferative wing discs of the *anchor* RNAi flies as not caused by apoptosis-induced compensatory proliferation. Hence, although *anchor* RNAi caused the transformation of interveins into more densely packed vein tissues, we also observed an increase in wing size, and these results demonstrate that the mechanism by which Anchor controls wing size is mainly mediated by the regulation of cell proliferation and cell size in imaginal discs.

### Knocking down *anchor* induced BMP downstream target genes in imaginal discs

The ectopic veins phenotype in the *anchor* RNAi wing disc was similar to that observed in gain-of-function of BMP signaling experiments, which were demonstrated by inducing the excessive phosphorylation of Mad (Mothers against dpp) (Haerry, 2010). Therefore, we next analyzed BMP signaling activity in knockdown *anchor* flies using p-Mad antibodies. We observed that p-Mad had a normal expression pattern along the AP boundary in *A9>anchor* RNAi discs but that its total intensity was clearly enhanced (Fig. 4A, B, K). Ectopically expressing either Dpp or excessively activating the Dpp receptor caused the overgrowth of the wing disc and increased the expression of its downstream targets (Nellen et al., 1996. Haerry et al., 1998). The markers *omb*, *sal*, *dad* and *brk* contain p-Mad dependent expression domains and are expressed along the AP boundary (Minami et al., 1999). Therefore, we analyzed the downstream transcriptional targets of p-Mad to identify the point of coupling between p-Mad and its targets in *anchor* knockdown flies. We showed that the brightness and widths of the *omb* and *sal* expression domains were strongly increased in *anchor* knockdown discs; however, *dad* and *brk* levels were similar to the levels observed in the control, and their total intensity was not changed (Fig. 4C-J, L-O). In pupal wings, *dad* levels were significantly increased; however, *brk* levels were reduced (Fig. S5). This result suggested that there is antagonism between Anchor and BMP signaling during wing development. However, the increased levels of p-Mad were not enough to induce changes in *dad* and *brk* levels in the imaginal disc.

**Fig. 4.**
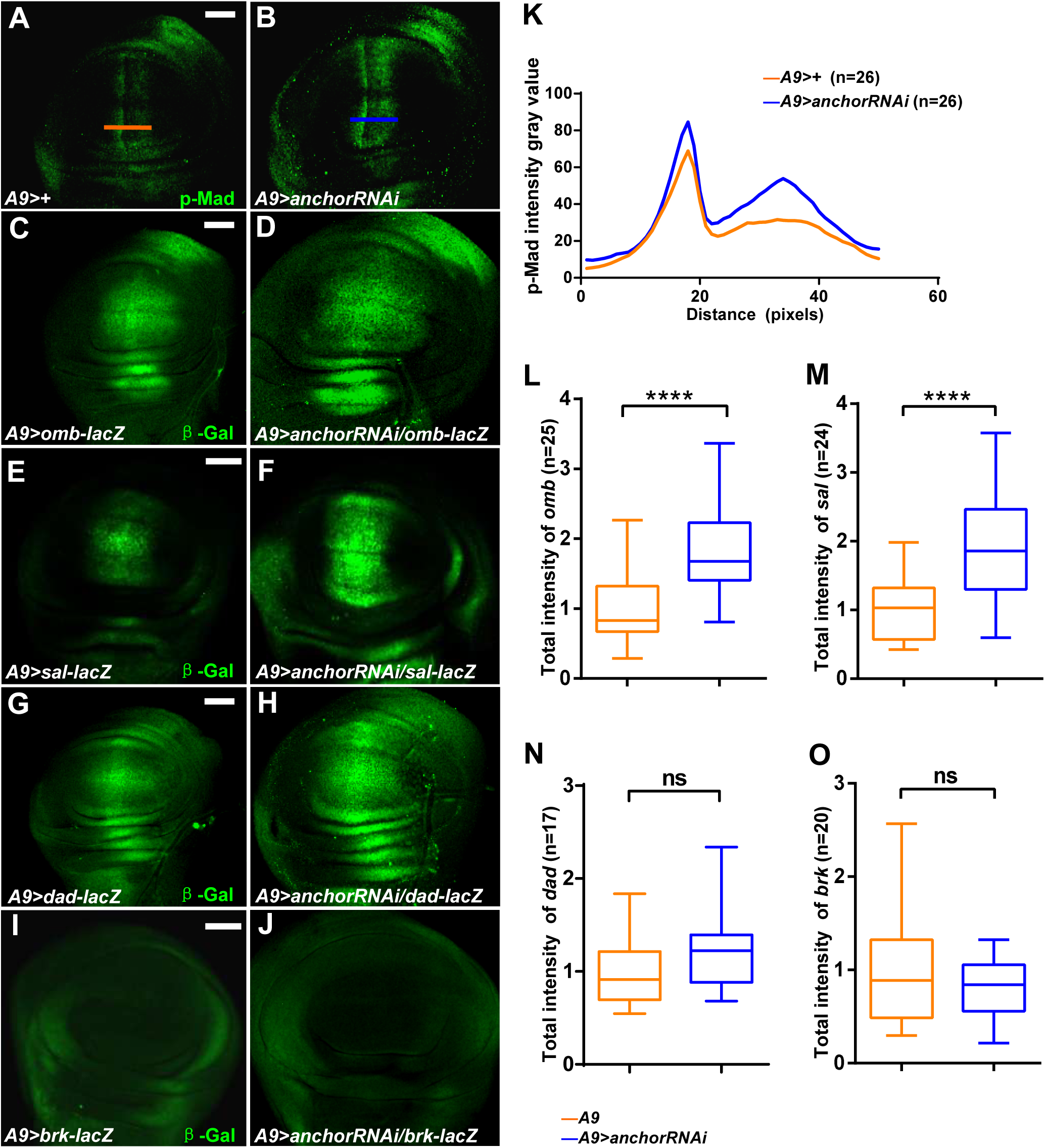
Analysis of BMP signaling pathway activity in *anchor* RNAi flies. (A-B, K) Immunohistochemical staining for p-Mad (green) in *anchor* RNAi flies showed that the p-Mad expression domain was larger in the mutant wing discs than in the control discs (A-B). The canonical late L3 p-Mad profile is outlined with blue and orange lines in *A9>anchor* RNAi and *A9>+* flies in the dorsal wing pouch, as shown in A and B (K). (C-J) The expression levels of BMP target genes, including *omb*, *sal*, *dad* and *brk*, were determined using *lacZ* reporter genes. The levels of *omb* and *sal* were significantly increased (C-F); however, *dad* and *brk* levels were not significant altered and were similar to the levels observed in the controls (G-J). (L-O) Quantification of the total intensity values of the four BMP target genes shown in C-J. All combinations were grown at 29°C, and female larvae were dissected. Scale bar: 50 μm.

### Anchor maintains the normal presumptive veins formation through limit BMP signaling in pupa stage

During metamorphosis, Dpp signaling is dramatically altered in the *Drosophila* wing. In the imaginal wing disc, the level of p-Mad reflects the gradient of Dpp, and this pattern is maintained during the early stages of wing metamorphosis. The p-Mad gradient is then lost and re-localized to presumptive veins (de Celis, 1997; O’Keefe et al., 2014) (Fig. 5A-A’, C-C’). Because of these shifts in the Dpp signaling zone, we hypothesized that *anchor* may regulate presumptive vein cell differentiation. We stained the intervein regions with anti-*Drosophila* Serum Response Factor (DSRF). We showed that p-Mad is roughly overexpressed, that its pattern is disordered in *anchor* knockdown pupal wings, and that edge veins excessively accumulated (Fig. 5B-B’, D-D’). This result indicates that p-Mad is also dramatic increased during the early pupal stage in *anchor* knockdown flies, which led to an ectopic vein phenotype in adult wings. Therefore, we concluded that *anchor* is essential for controlling BMP signaling to safeguard the proper formation of normal wing veins during early pupal stages.

**Fig. 5.**
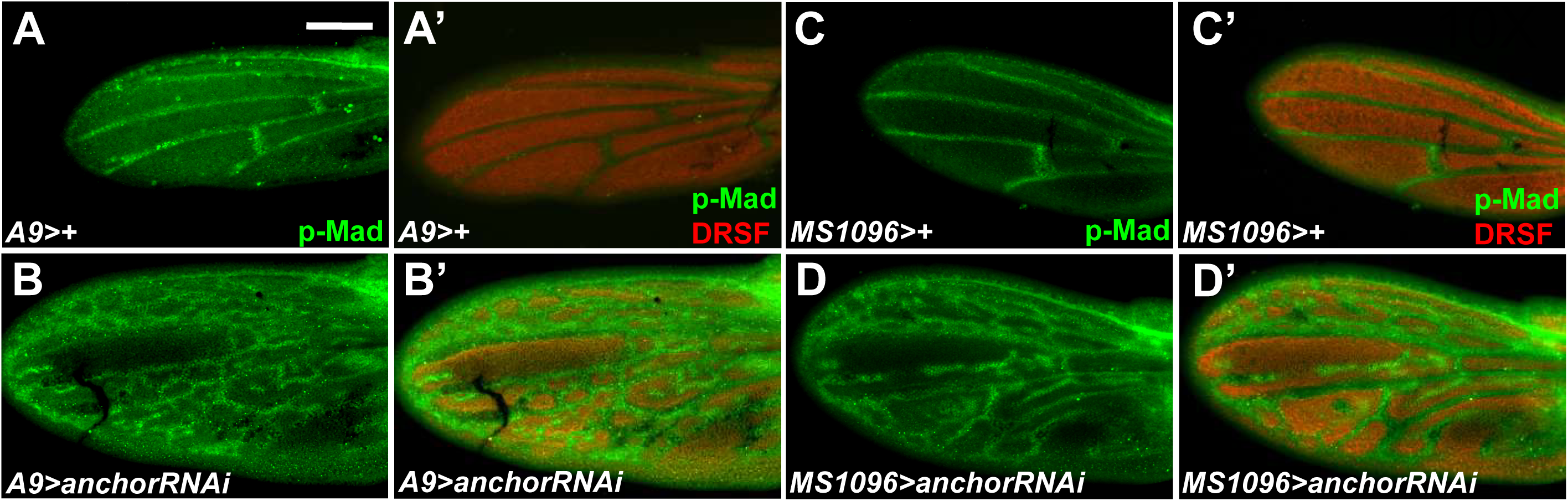
Increased and ectopic accumulation of p-Mad in presumptive veins and at the margin in *anchor* RNAi flies during pupal development. Immunochemical staining showing the wing blade and interveins was performed using p-Mad (green) and DRSF (red) antibodies at 24 hr APF. p-Mad was localized to presumptive veins and the anterior margin in control pupal wings (A-A’ C-C’). After *anchor* knockdown, pupal wings displayed a massive amount of ectopic p-Mad, and it accumulated in cells near the distal wing margin (B-B’, D-D’). All combinations were grown at 29°C. Scale bar: 100 μm.

### Anchor targets *dpp* or works upstream of *dpp* to antagonize BMP signaling

During the pupal stage, Dpp signaling is critically required for wing vein cell differentiation and for these cells to acquire the appropriate location (de Celis, 1997). Ectopically expressing either *dpp* or activating Dpp signaling resulted in thickened or extra veins (O’Keefe et al., 2014). To further investigate the relationships between Anchor and Dpp signaling, we used a genetic approach to perform a rescue experiment. Downregulating the Dpp signaling mediators *med* and *mad* using RNAi in flies induced the removal of parts of L3, L4 and L5 (Fig. 6G-H). However, combining the depletion of *med* or *mad* with *A9>anchor* RNAi flies limited the width of veins (Fig. 6A, M-N). Similarly, knocking down *anchor* using the background null allele *mad*^*12*^ also rescued the broadening of vein tissues (Fig. 6B-C). This result indicated that *anchor* is required to limit vein tissue formation, while *med* or *mad* are downstream effectors of *anchor*. We also investigated the genetic relationship between *anchor* and Tkv, Sax or Put receptors, which are involved in BMP signaling. The hypomorphic alleles *tkv*^*7*^, *tkv*^*427*^ and *sax*^*4*^ or the RNAi flies *A9>sax* RNAi and *A9>put* RNAi resulted in significantly suppressed thickened vein tissue that was caused by a reduction in *anchor* levels (Fig. 6A-B, D-F, I-J, O-P). The *mad*^*12*^/+, *sax*^*4*^/+, *tkv*^*7*^/+, and *tkv*^*427*^/+ flies exhibited normal adult wing phenotypes (data not shown).

**Fig. 6.**
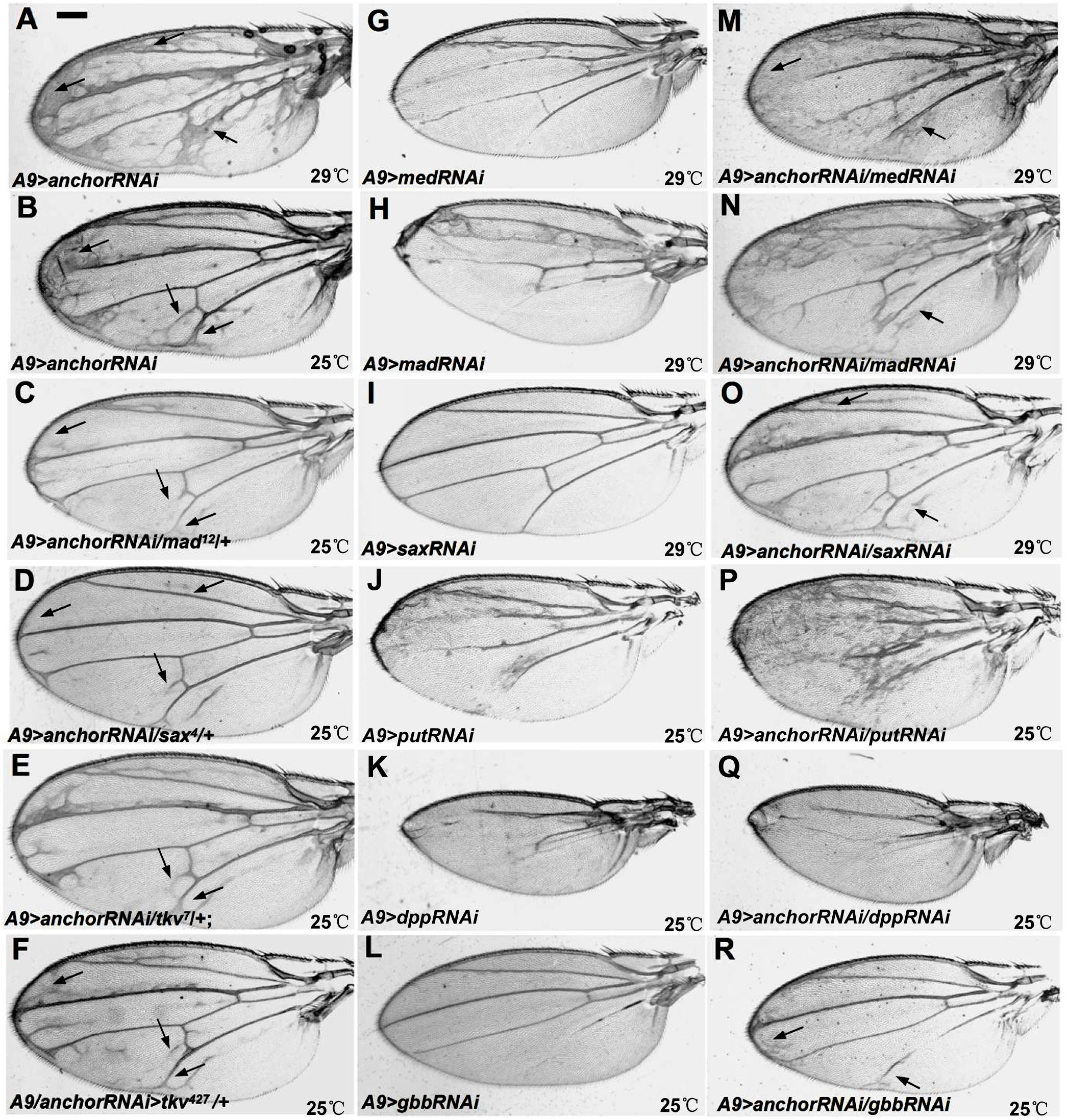
Anchor antagonizes the BMP signaling pathway. (A, B) *A9>anchor* RNAi resulted in broadened L3, L5 and distal marginal veins and ectopic L5 and PCV veins, as indicated by arrows. (C) *A9>anchor* RNAi/*mad*^*12*^ suppressed the development of distal marginal veins and broadened L5 (C versus B, arrows). (D) *A9>anchor* RNAi/*sax*^*4*^ suppressed the broadening of distal marginal veins (D versus B, arrows). (E) *A9>anchor* RNAi/*tkv*^*7*^ suppressed the broadening of L5 (E versus B, arrow). (F) *A9>anchor* RNAi/*tkv*^*427*^ suppressed the broadening of distal marginal veins and L5 (F versus B, arrow). (M) *A9>anchor* RNAi/*med* RNAi suppressed the broadening of distal marginal veins (M versus A, arrow), and the low expression of *med* resulted in shortened L3, L4 and L5 (M versus G). (N) *A9>anchor* RNAi/-*mad* RNAi suppressed the broadening of L5 (N versus A, arrow), and the low expression of *mad* resulted in shortened L3, L4 and L5 (N versus H). (O) *A9>anchor* RNAi/*sax* RNAi suppressed the broadening of L2 (O versus A, arrow). (P) *A9>anchor* RNAi/*put* RNAi resulted in a phenotype similar to that of the *put* RNAi (P versus J), and the missing PCV and missing portion of L3-5 resulted from the low expression of *put*. (Q) *A9>anchor* RNAi/*dpp* RNAi resulted in a phenotype similar phenotype to that of the *dpp*RNAi (Q versus K), and the missing veins resulted from the low expression of *dpp*. (R) *A9>anchor* RNAi/*gbb* RNAi suppressed the broadening of L3, L5 and distal marginal veins (R versus A), and the missing PCV and the missing portion of L5 was caused by the low expression of *gbb* (R versus L). Other details of these experiments are analyzed in Table S1. Scale bar: 200 μm.

In addition, knocking down *dpp* or *gbb* severely depleted vein tissue and reduced wing size (Fig. 6K-L). However, knocking down *dpp* in *A9>anchor* RNAi flies did not rescue the loss of wing vein phenotype, and the adult wing veins displayed a phenotype that was similar to the phenotype caused by the *A9>dpp* RNAi (Fig. 6 K-Q). In addition, the reduced wing size and loss of veins that were caused by *gbb* RNAi were rescued when the RNAi was combined with *A9>anchor* RNAi flies, although the distal veins remained slightly thickened (Fig. 6B, L-R). This result indicates that Dpp is required for the ectopic vein formation that is caused by knocking down *anchor*. Thus, the *dpp* depletion phenotype was epistatic to the reduction in *anchor*, which shows that Dpp is required for the reduction in wing veins that was observed in the *anchor* phenotype. The genetic epistasis test suggested that *anchor* is likely to function upstream of Dpp signaling or by targeting Dpp through a parallel pathway.

## DISCUSSION

In this study, we have analyzed the function of novel gene *anchor* in wing development. Inducing the wing-specific knockdown of *anchor* resulted in the formation of thickened and ectopic veins. Moreover, this phenotype was similar to that observed in gain-of-function BMP signaling experiments. In addition, we also observed enlarged wing blades and an increase in wing size in the *anchor* RNAi flies because of the high number of PH3-positive cells and enlarged wing blade cells. In addition, the intensity profile for p-Mad was enhanced in *anchor* RNAi-induced imaginal discs. Hence, Anchor substantially contributed to limiting the phosphorylation of Mad in presumptive veins during the pupal stage, and accordingly, the BMP signaling targets genes were significantly up-regulated. In genetic interaction experiments, we found that *anchor* was most likely functioning upstream of *dpp* or that it targeted *dpp*. We concluded that Anchor functions during wing vein development to inhibit BMP signaling.

### Anchor regulates cell proliferation in wing discs

When the *anchor* RNAi was expressed throughout the wing disc in *Gal4* fly lines, it caused an increase in wing size in larvae and adults. Many different types of signals participate in cell proliferation within developing tissues. For example, Wg and Dpp signals mediate wing disc growth in different ways. The morphogen Wg is expressed at the wing pouch boundary to delineate the domain of the wing blade within the wing disc. The involvement of Wg signaling in compensatory proliferation has been explored by examining its expression in apoptotic cells. In apoptotic cells, ectopic Wg signaling induced compensatory proliferation during mitosis (Martín et al., 2009). However, when we used TUNEL assays, the number of apoptotic cells was not changed when Wg intensity was increased in *anchor* RNAi-induced wing discs. Therefore, increased Wg intensity was another factor that enlarged wing blades in the *anchor* RNAi flies. The morphogen Dpp is expressed along the AP boundary, where it forms a gradient during wing disc development. Dpp signaling is thought to promote growth and proliferation. When *dpp* was overexpresses or enhanced in the wing disc, it led to overgrowth or enlarged wing size in adult flies (Martín-Castellanos and Edgar, 2002; O’Keefe et al., 2014). We observed a dramatic increase in the levels of p-Mad, *sal* and *omb* in the *anchor* knockdown flies, which indicated that Anchor opposed Dpp signaling to inhibit cell proliferation in wing discs.

### Dpp signaling rebuilds after pupariation

In wing imaginal discs, Dpp is expressed in a strip of cells close to the anterior edge of the AP boundary. It transmits its signal via receptor complexes and transducer dimers (Israel et al., 1996). After pupariation, Dpp signaling continues to be expressed at the AP boundary and also appears along presumptive veins. This shift in expression results in subsequent changes in p-Mad and downstream targets. At 24 hr APF, p-Mad expression is restricted to presumptive and marginal vein cells. *omb* is expressed in the distal edge of the pupal wing, and *dad* expression is broadly activated in cells corresponding to presumptive veins. At the same time, the expression of *brk* is eliminated from presumptive veins and becomes restricted to intervein cells (Sotillos and De Celis, 2005). p-Mad levels are substantially increased in pupal wings that express low levels of *anchor* at 24 hr APF. Its transcriptional targets, including *omb* and *dad*, are also substantially increased (Fig. S3). Consistent with p-Mad expansion in intervein region, brk expression is abolished from essentially intervein cells in *anchor* RNAi pupal wing. These results are evidence that *anchor* is involved in Dpp signaling and that it maintains the position of wing veins and distinguishes the boundaries between veins and interveins.

### *anchor* interactions with ligands antagonize BMP

Our experiments reveal the following important factors that contribute to the connection between Anchor and BMP signaling: (1) decreasing *anchor* levels in the wing discs induces the overexpression of BMP signaling factors, which increases wing blade size and adult wing size; (2) in these flies, the levels of BMP signaling effectors, including p-Mad, *sal*, and *omb*, are generally increased in the wing discs, with p-Mad levels being especially increased during the pupal stage; and (3) mutations in BMP signaling factors rescued the ectopic phenotypes observed in *A9>anchor* RNAi-induced flies to varying degrees. These data indicate that in the absence of *anchor*, BMP signaling is enhanced.

In normal wing disc cells, the Dpp ligand prefers to bind to Tkv, while the Gbb ligand has a higher affinity for Sax. The type I receptor Tkv is required for the transcription of all BMP signaling target genes during wing development, and Sax promotes BMP signaling by forming a receptor complex with Tkv. In wing discs, Tkv and Sax can form three types of receptor dimer complexes: Tkv-Tkv, Tkv-Sax and Sax-Sax. The Tkv-Sax receptor complexes are probably the mechanism by which Sax contributes to promoting BMP signaling, which initiates a more robust intracellular phosphorylation cascade than the Tkv-Tkv dimers (Bangi and Wharton, 2006). However, Sax-Sax dimers initiate almost no signaling (Weis-Garcia and Massagué, 1996; Haerry, 2010). The Tkv-Sax receptor complexes result in more substantial signaling than the other dimer complexes. Anchor appears to preferentially bind to the ligand Dpp (Fig. 7E). Knocking down *anchor* induced p-Mad levels to increase, which caused adult wings to exhibit thickened and ectopic veins. Anchor was not available to control the ligand Dpp, which could then induce the increased phosphorylation of Tkv homodimers (Fig. 7A and F). We found that overexpressing *tkv* or *dpp* enhanced the extent of ectopic vein tissue to background *anchor* knockdown levels (Fig. 7B and C). Compared to the effects of the *anchor* RNAi, overexpressing *tkv* or *dpp* increased the number of Tkv receptor homodimers, which led to a moderate increase in phosphorylation (Fig. 7G). Substantially overexpressing *gbb* allowed Dpp and Gbb to activate Tkv-Sax heterodimers to undergo high levels of phosphorylation in *anchor* deletion mutants (Fig. 7D and H). Therefore, we hypothesized that Anchor restricts the ligand Dpp from inducing the excessive formation of heterodimers or homodimers, which led to a large increase in p-Mad levels.

**Fig. 7.**
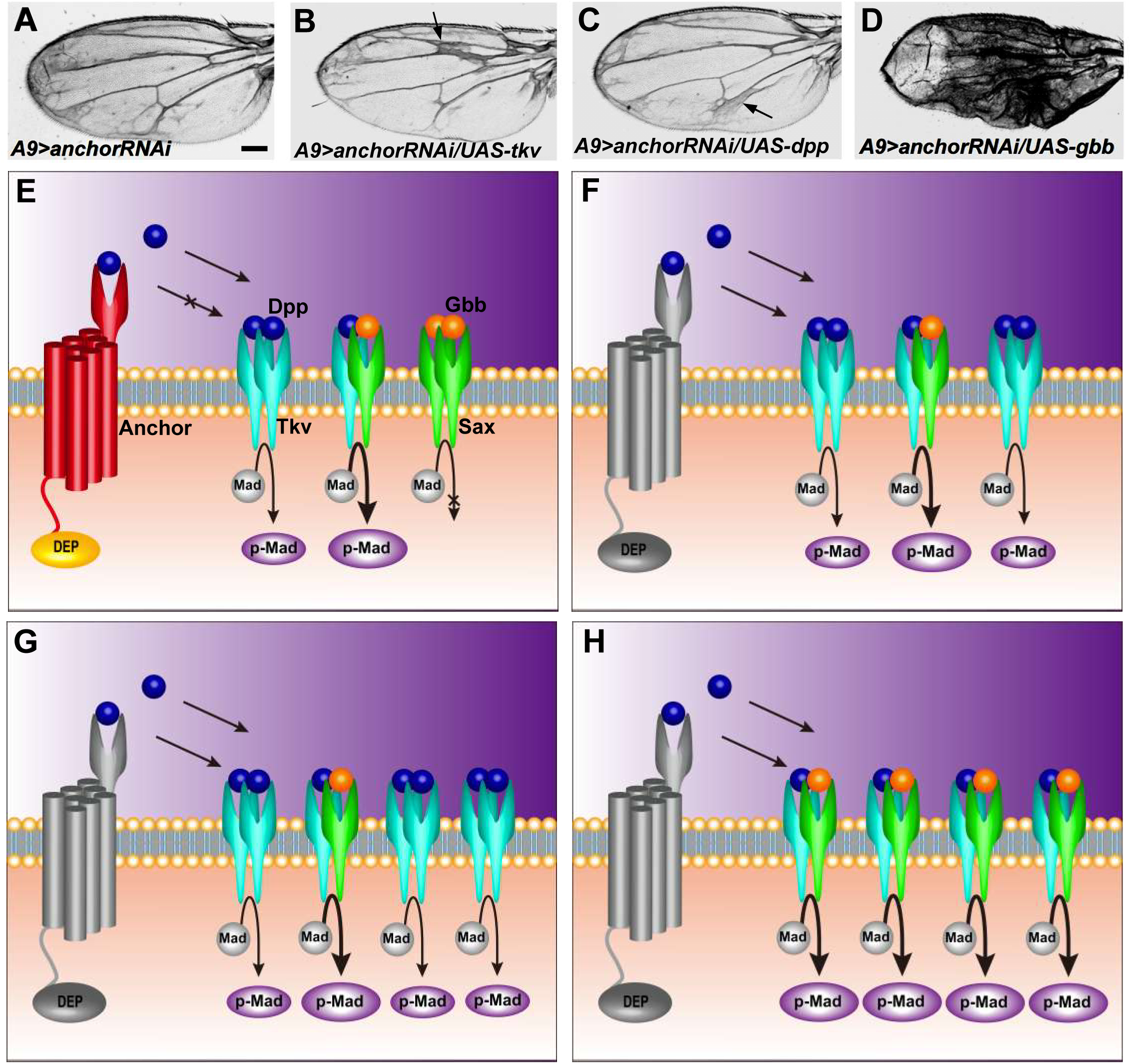
Hypothetical Models explaining the opposing effects of Anchor on the BMP signaling pathway. (A-D) Inducing the overexpression of *tkv* or *dpp* (BMP effectors) in *anchor* RNAi flies enhanced the thickened and ectopic vein phenotype (B and C, arrows). Similar to the effect of overexpressing *gbb*, these flies formed massively ectopic veins (D). All of the flies were grown at 25°C, scale bar: 200 μm. (E) A model showing both the antagonistic functions and the signaling functions of *anchor* in wing disc cells. The ligands displayed preferential affinity to different receptors, with Dpp preferentially binding to Tkv receptors and Gbb preferentially binding to Sax receptors. The different receptor complexes induced varying degrees of phosphorylation degrees. Tkv-Sax receptor complexes mediated more significant levels of signaling (thickened arrow) than were initiated by Tkv-Tkv complexes (thin arrow). However, Sax-Sax complexes did not phosphorylate Mad. Anchor restricts the free ligand Dpp from being released from the intracellular compartment, to some extent. (F) Cells expressing low levels of *anchor* failed to limit the ligand Dpp, resulting in more Dpp binding to each to increase the levels of p-Mad. (G) Overexpressing *dpp* or tkv in *anchor* RNAi cells contributed to a higher level of p-Mad than was observed in *anchor* RNAi cells (compare C to A). (H) In the absence of *anchor*, overexpressing *gbb* led to the ligands inducing the formation of heterodimers. These dimers bind to Tkv-Sax receptor complexes to induce the levels of p-Mad to increase (compare D to A).

When we simultaneously reduced *anchor* and *gbb* expression, we observed adult wing veins that displayed normal longitude veins but distal veins that were slightly broadened (Fig.6B, L, R). This phenomenon indicated that *anchor* affected Gbb signaling but did not directly interact with the ligand Gbb. When we simultaneously reduced *anchor* and *dpp* levels, we observed adult wing veins that displayed a phenotype like that of the *dpp* RNAi-induced flies, which lost almost all of their veins (Fig. 6B, K and Q). These data further confirm that *anchor* is likely to function either upstream of Dpp signaling or through a parallel pathway. We propose the hypothesis that Anchor traps the Dpp ligand to negatively regulate the BMP signaling pathway during wing development.

## METERIALSAND METHODS

### Fly stocks

The two *anchor* RNAi forms (v8532, v105969) were obtained from the Vienna *Drosophila* RNAi Stock Center (VDRC). The *med* RNAi, *mad* RNAi, *dpp* RNAi, *gbb* RNAi, *sax* RNAi, and *put* RNAi were obtained from Tsinghua *Drosophila* model animal center. The *dpp-Gal4*, *ap-Gal4*, *patched-Gal4*, *brk-Gal4*, *en-Gal4*, *omb-lacZ*, *dad-lacZ*, *brk-lacZ*, null allele *Mad*^*12*^, *tkv*^*427*^, *tkv*^*7*^ and *sax*^*4*^ flies were described previously (Liu et al., 2011). The *sal-lacZ* was generously provided by J Shen (University of China Agriculture). The *UAS-anchor* was obtained by germline transformation. A strain in which *anchor*^*G9098*^ was inserted using *P* elements into the 5’ region of the *anchor* gene was purchased from GenExel (Daejeon, Korea). Other strains used in this study included *UAS-dpp*^*19B5*^, *UAS-gbb*^*99A2*^, *UAS-tkv*^*1A3*^, *A9-Gal4*, *MS1096-Gal4*, *ptc-Gal4*, and *w*^*1118*^. All genotypes were bred into the *w*^*1118*^ background. In this study, we used another *anchor* RNAi construct (v105969) to knockdown *anchor* levels, as shown in Fig. S2.

### *In Situ* Hybridization

The partial sequence of *anchor* (1-600 bp) was amplified from cDNA using PCR and subcloned into the pSPT18/19 vector. Digoxigenin-labeled riboprobes were prepared using T7 RNA polymerase and a DIG RNA labeling kit (Roche). RNA *in situ* hybridization on third instar larval (L3) wing discs was carried out according to a standard protocol. The tissue was fixed in RNAse-free 4% paraformaldehyde in PBS for 1 hr. The wing discs were prehybridized with 100 μg of salmon sperm DNA/ml in hybridization buffer at 55°C for 1 hr. Hybridizations were performed overnight at 55°C in hybridization solution containing an *anchor* probe. The wing discs were incubated with alkaline phosphatase-conjugated anti-DIG antibodies (Roche) for 2 hr at room temperature, and hybridization signals were visualized using BCIP/NBT. Finally, the slides were mounted using Vectashield mounting medium (Vector Laboratories) and analyzed using an Axioskop 2 plus microscope (Zeiss). All experiments were independently repeated at least three times

### Immunohistochemistry

Wing imaginal discs obtained from third instar larvae were fixed in 4% paraformaldehyde for 1 hr at room temperature. The wing discs were then placed in blocking buffer (PBS plus 0.1% Tween 20 and 5% normal goat serum) for 1 hr at room temperature. To prepare pupal wings, late third instar larvae were selected and allowed to develop for 24 hr after puparium formation (APF) at 29°C. Whole pupae (24 hr APF) were removed from their pupal cases and fixed in 3.7% formaldehyde for 2 hr at room temperature. The pupal wings were dissected in 0.3% PBST (0.3% TritonX-100 in PBS), incubated in blocking buffer (PBS containing 0.3% Triton X-100, 2% BSA, and 2% normal goat serum) for 1 hr (Liu et al., 2011). The wings were incubated in primary antibodies overnight at 4°C and then incubated with secondary antibodies according to standard methods. Finally, the wings were mounted in Vectashield fluorescent mounting medium (Vector Laboratories) or Prolong diamond antifade mountant (Molecular probes). The tissues were analyzed using a LSM 510 META confocal microscope (Zeiss) or an Axioskop 2 plus microscope (Zeiss). The following primary antibodies were used: mouse anti-β-gal (1:200, Promega), rabbit anti-pSmad1 (1:50, a gift from Ed Laufer, Columbia University, New York) (Vargesson and Laufer, 2009), rabbit phospho-H3 (1:800, Upstate), and mouse anti-DSRF (1:100, Active Motif). Secondary antibodies were conjugated with Alexa Fluor 488 and Alexa Fluor 568 (Molecular probes) and used at a 1:200 dilution. All experiments were independently repeated at least three times.

### Transgenic constructs

To generate a *UAS-anchor* strain, the full-length *anchor* coding sequence was PCR amplified and cloned into pUAST between the underlined restriction sites. The following primers were used: 5′-AAAGAATTCATGGACAGCTCCATGTACTACG-3′ and 5′-AAACTCGAGCTATATGCGACTGCAGAAAT-3′. Transgenic flies were then generated using standard methods.

### Statistical Analysis

Images were acquired using a Zeiss fluorescence microscope. All numerical data, including wing size, imaginal disc size, cell size, cell number and intensity values, were analyzed using image J. The statistical analyses were performed using a two-tailed unpaired Student’s *t*-test with Prism software (GraphPad 6.0). \*\**P*<0.005 was considered significant, and \*\*\**P*<0.001 was considered more significant. “ns” indicates no significant difference. Error bars in the graph indicate SEM.

## Acknowledgements

We are grateful to Ed Laufer or generously providing anti-pSmad1. We thank J Shen for supplying us with the strian *sal-lacZ*. We gratefully acknowledge Vienna *Drosophila* RNAi Stock Center, Tsinghua *Drosophila* model animal center, GenExel Stock Center and Developmental Studies Hybridoma Bank for providing fly lines and antibodies.

## Competing interests

The authors declare no competing or financial interests.

## Funding

This work was supported by the National Natural Science Foundation of China (31270923) and the Fundamental Research Funds for the Central Universities China (2572015AA10).

---

**Fig. S1.**
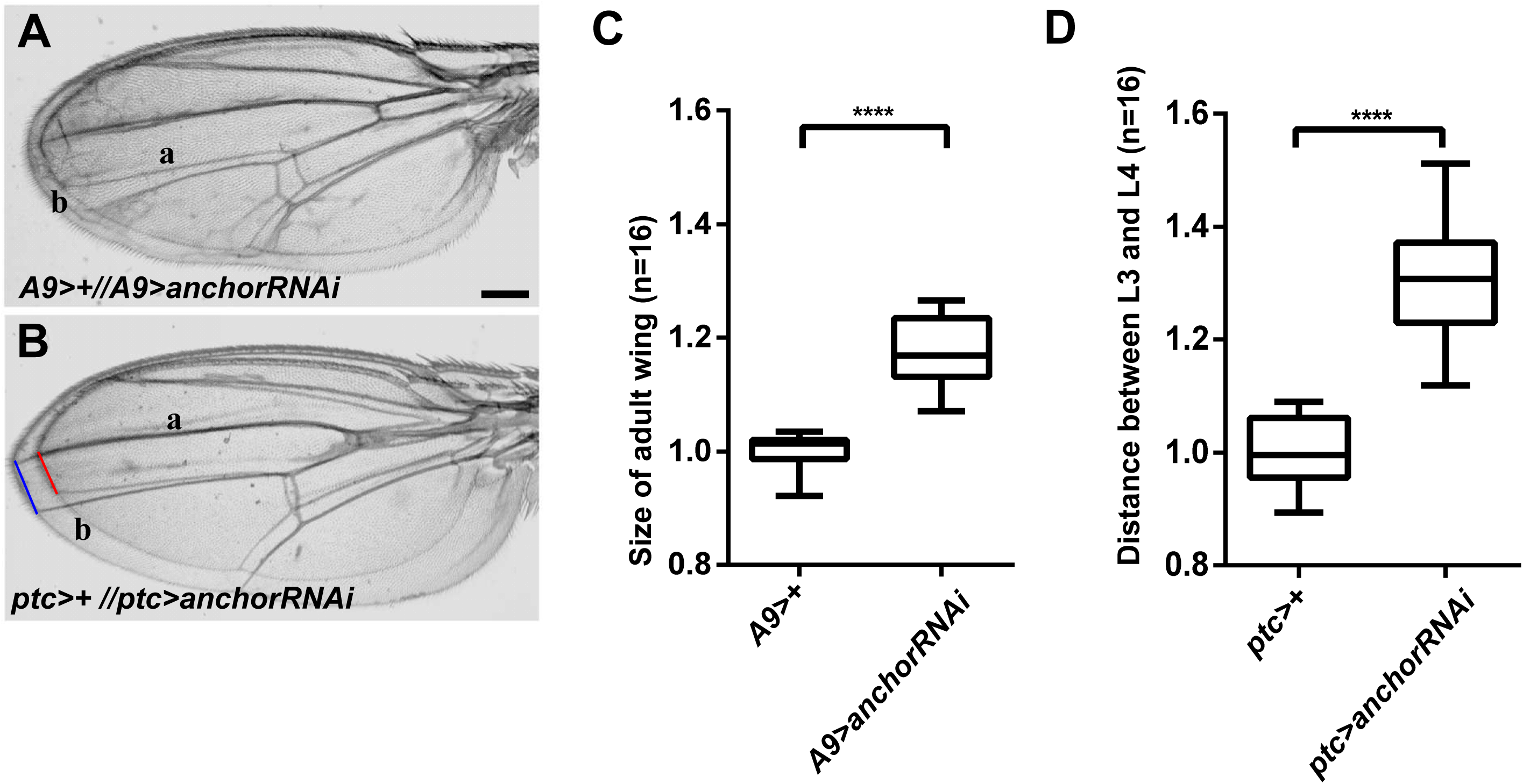
Wing-specific knockdown of *anchor* caused enlarged wing size in adult flies. (A) Overlapped adult wings obtained from *A9>+* and *A9>anchor* RNAi flies show that wing size was larger in the anchor RNAi flies. (B) Overlapped adult wings obtained from *ptc>+ and ptc>anchor* RNAi flies show that the distance between L3 and L4 was longer in the *anchor* RNAi wings. (C, D) The quantification of wing size and the distance between L3 and L4 are compared in A and B. All flies were grown at 25°C, scale bar: 200 μm.

**Fig. S2.**
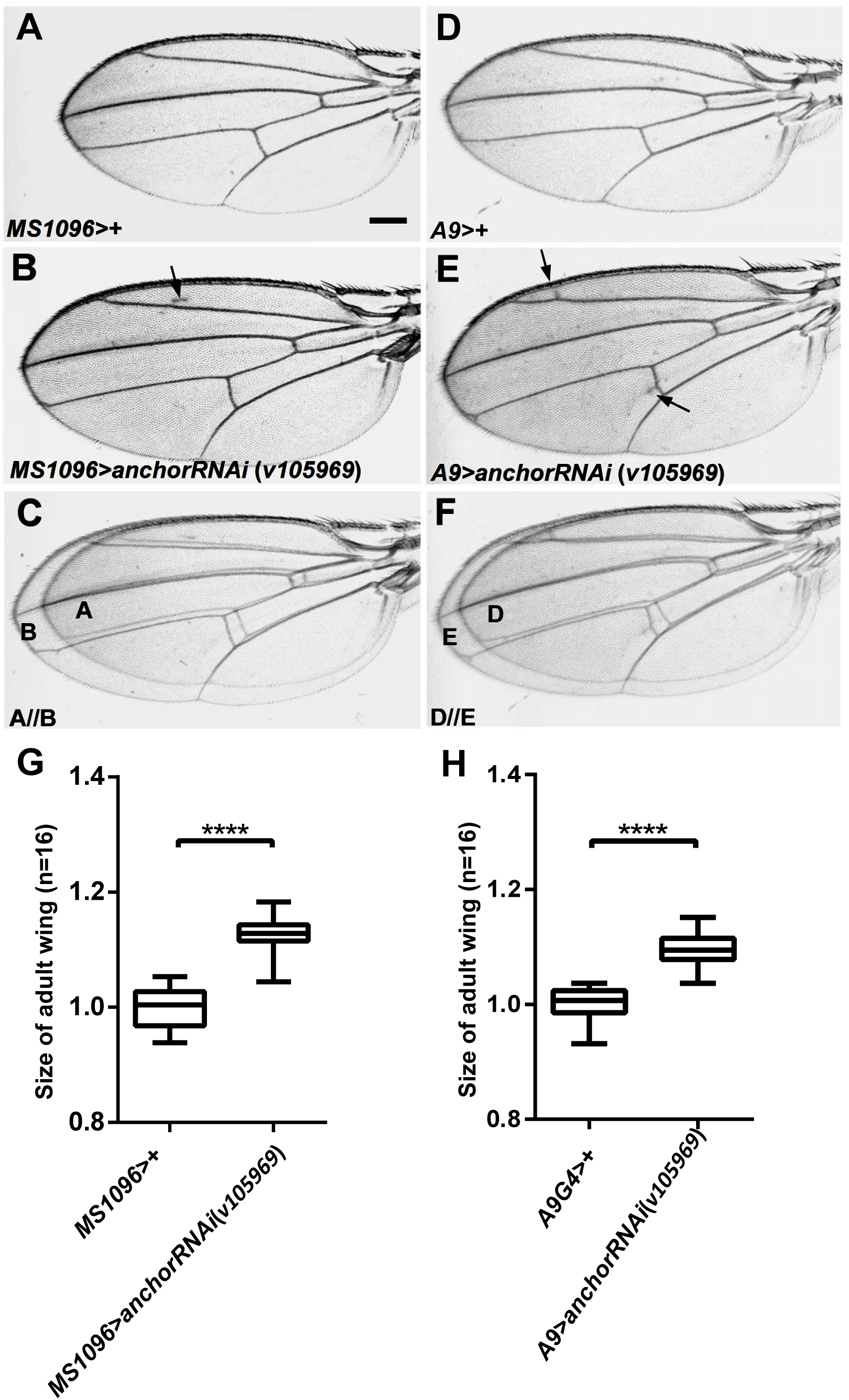
Knockdown of *anchor* using *anchor* RNAi (v105969) resulted in a phenotype similar to that of *anchor* RNAi (v8532) flies, which displayed extra veins and larger wings. (A-F) Knocking down *anchor* using *MS1096>anchor* RNAi (v105969) and *A9>anchor* RNAi (v105969) resulted in extra veins and enlarged wings, similar to the anchor RNAi (v8532) phenotype. (G, H) The quantification of wing size, as compared in C and F. All flies were grown at 29°C, scale bar: 200 μm.

**Fig. S3.**
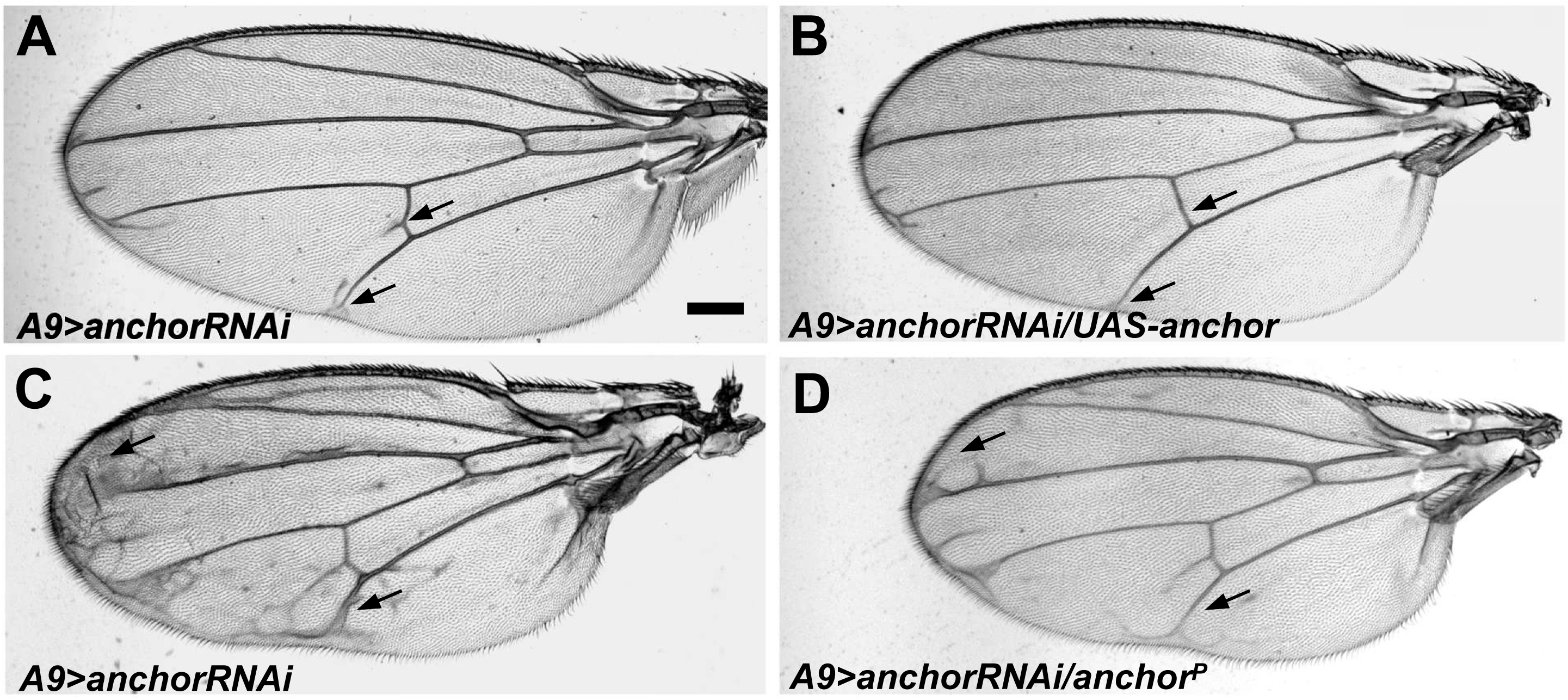
Overexpressing *anchor* rescued the ectopic phenotypes in *anchor* RNAi flies. Overexpressing full length *anchor* using *UAS-anchor* or *anchor^P^* in RNAi flies significantly suppressed extra vein formation in adult flies (B compared to A). The *anchor*^*P*^ flies contained a *P*-element insertion in the 5’ UTR region of *anchor* that encoded a Gal4-responsive enhancer (D compared to C). All crosses were performed at 18°C (A-B) or 25°C (C-D). Scale bar: 200 μm.

**Fig. S4.**
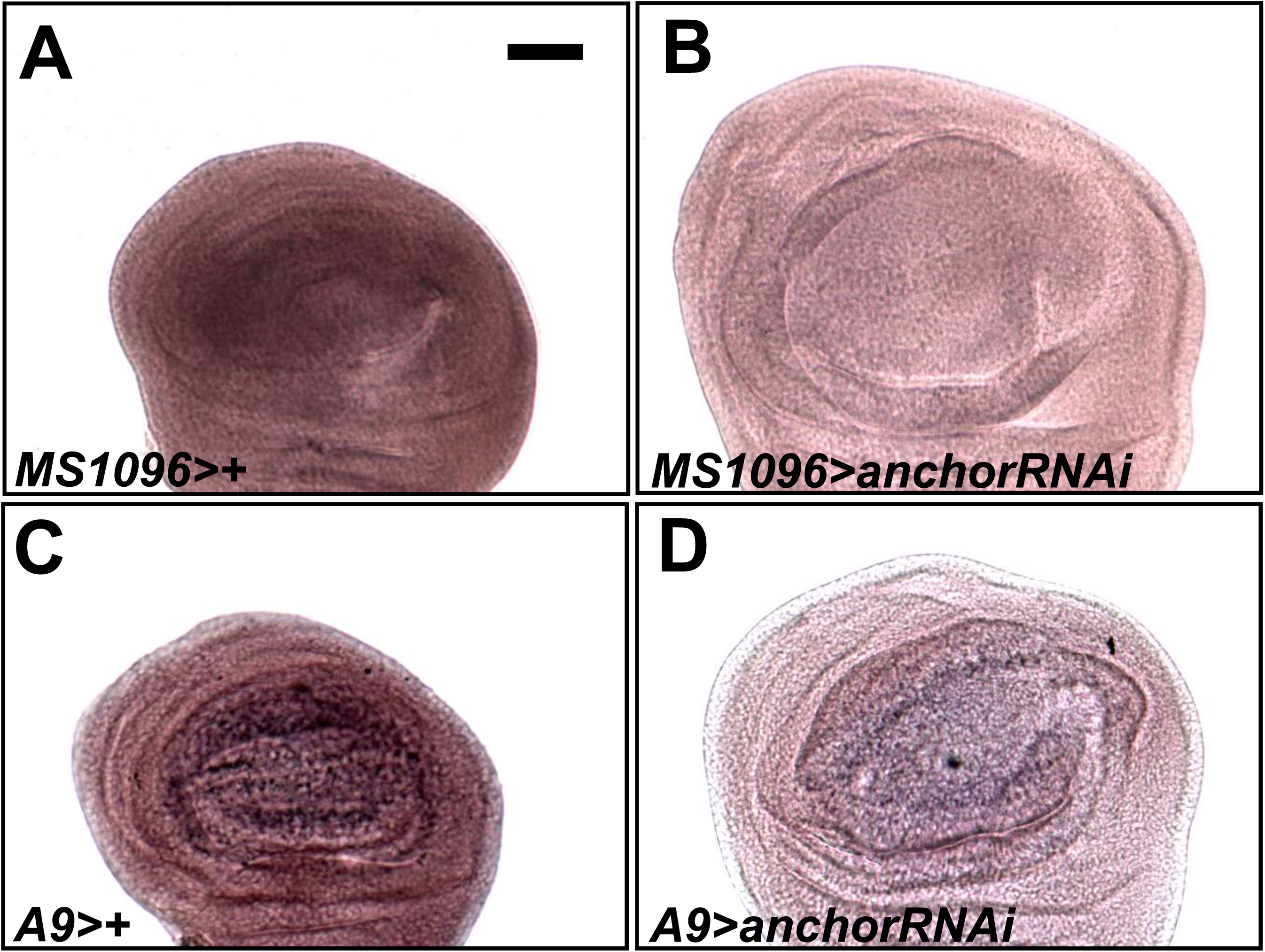
Distribution of *anchor* gene expression in wing discs. *In situ* hybridization was performed using an *anchor*-specific probe in L3 wing discs. (A, C) In control discs, the *anchor* gene is expressed in a generalized manner. (B, D) When an *anchor* RNAi driven by *MS1096-Gal4* or *A9-Gal4* was used, we observed a significant reduction in *anchor* gene levels. In addition, *anchor* RNAi caused the wing blade region to overgrown within the imaginal disc. All combinations were grown at 29°C, and selected female larvae were dissected. Scale bar: 50 μm.

**Fig. S5.**
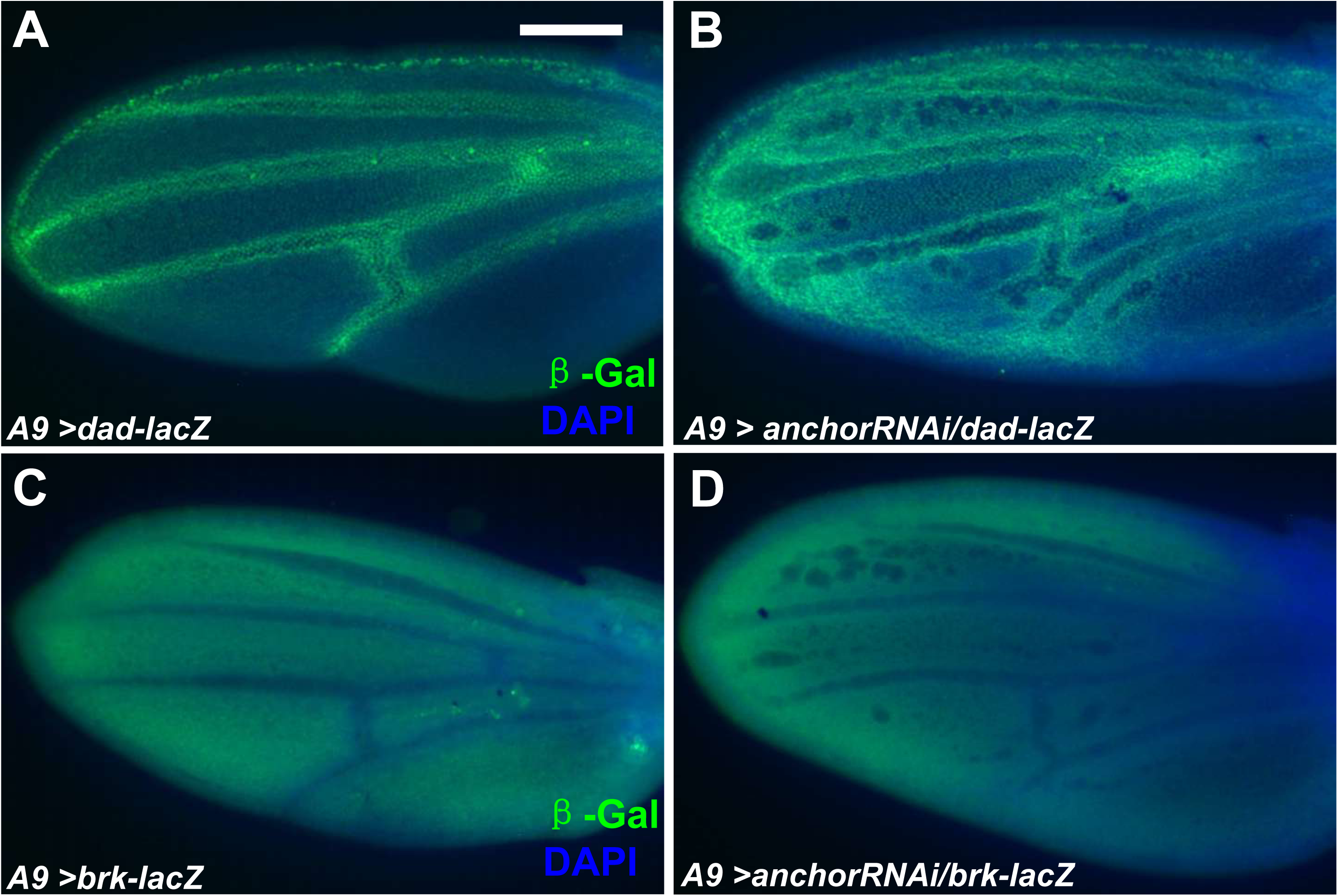
Knocking down *anchor* shifted *dad* and *brk* levels in pupal larvae. The expression levels of the BMP targets *dad*and *brk* were assayed using *lacZ* reporter genes. The *dad* expression domain was significantly expanded in presumptive veins, and *brk* expression was reduced. All combinations were grown at 29°C, scale bar: 100 μm.

**Table S1.**
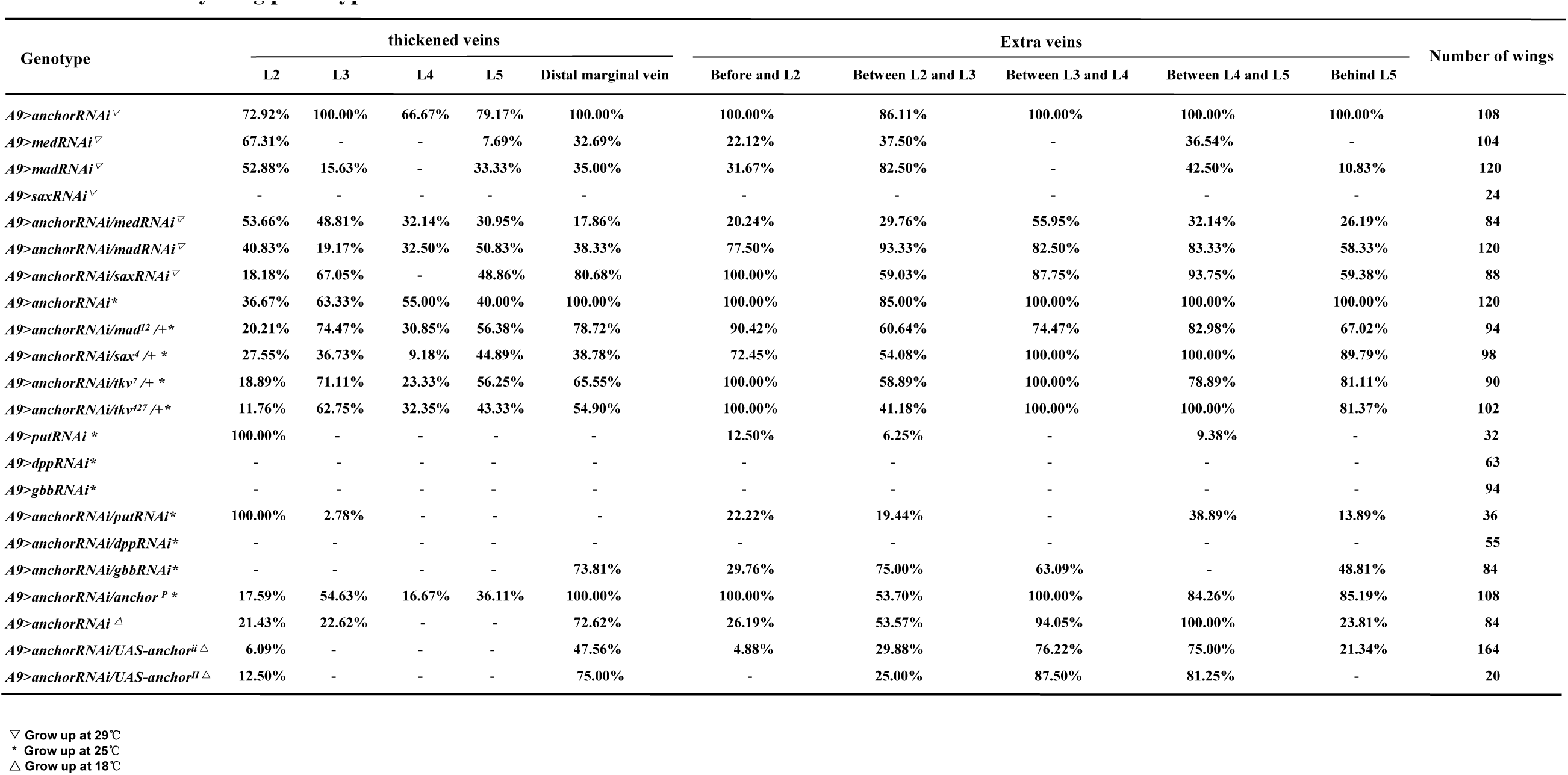
Summary wing phenotypes of *anchor* RNAi combinations

